# ArdC protein overpasses the recipient *hsdRMS* restriction system broadening conjugation host range

**DOI:** 10.1101/756429

**Authors:** Lorena González-Montes, Irene del Campo, Fernando de la Cruz, Gabriel Moncalian

## Abstract

Plasmids, when transferred by conjugation, must overpass restriction-modification systems of the recipient cell. We demonstrate that protein ArdC, encoded by broad host range plasmid R388, was required for conjugation from *Escherichia coli* to *Pseudomonas putida*, but not from *E. coli* to *E. coli.* Surprisingly, expression of *ardC* was required in the recipient cells, but not in the donor cells. Besides, *ardC* was not required for conjugation if the *hsdRMS* system was deleted in *P. putida* recipient cells. Thus, ArdC has antirestriction activity against HsdRMS system, and consequently broadens R388 plasmid host range. The crystal structure of ArdC was solved both in the absence and in the presence of Mn^2+^. ArdC is composed of a non-specific ssDNA binding N-terminal domain and a C-terminal metalloprotease domain, although the metalloprotease activity is not needed for antirestriction function. We also observed by RNA-seq that ArdC-dependent conjugation triggers an SOS response in the *P. putida* recipient cells. Our findings give new insights, and open new questions, into the antirestriction strategies developed by plasmids to counteract bacterial restriction strategies.

## Introduction

Horizontal gene transfer (HGT) is the transmission of genetic material between organisms that are not in a parent– progeny relationship [1]. The clinical relevance of the HGT process lies on the acquisition and dissemination of genes involved in conferring bacterial resistance to antibiotics (Ab^R^) between unrelated pathogens. When bacteria face selective pressures, as those exerted by antibiotics, horizontal acquisition of Ab^R^ allows diversification of the genomes, increasing survival opportunities. Conjugation is the main HGT process that allows the transfer of genes encoded in autonomous plasmids. This process requires the machinery to build a direct contact between a donor and a recipient cell [1]. Conjugation can be modulated by environmental factors or bacterial strategies based on genetic approaches that are coded in the chromosome (host barriers) or plasmid DNA (plasmid barriers). Plasmid barriers include entry exclusion [2] or fertility inhibitor [3] which reduce conjugative transfer. Host barriers can be mediated trough SOS response modulation [4,5], CRISPR-Cas systems [6] or restriction and modification (R-M) systems. R-M systems allow bacteria to discern between self-DNA and foreign DNA invading the cell, leading to its destruction. They require two enzymatic activities: a methyltransferase that provides protection to its own DNA and an endonuclease that cleaves the unmethylated invading DNA [7]. There are four main groups of R-M systems. Type I R-M system requires three genes: *hsdR*, *hsdM* and *hsdS* and their products associate in R_2_M_2_S complexes. S subunit recognizes 13-15 bp sequences, usually asymmetric and bipartite. DNA cleavage is at a location away from the specificity site [8–10]. There is a coevolutionary arms race between bacteria to avoid entrance of foreign DNA molecules and parasitic DNA molecules as plasmids or bacteriophages to enter a putative host avoiding the restriction by bacterial R-M systems. The antirestriction mechanisms to counteract R-M systems can be divided in four main types based on its mode of action: DNA modification, transient occlusion of restriction sites, sabotage of host R-M activities, and inhibition of restriction enzymes [9]. R388 plasmid is the prototype of the IncW incompatibility group of plasmids. IncW plasmids have a low copy number, a wide range of Ab^R^, and a broad host range (BHR) [11]. R388 has 35 genes assorted in functional groups or modules, between them, a gene coding for an antirestriction protein called ArdC [11]. Here, we present ArdC crystal structures and its role in interspecies conjugation. We have also identified transcriptional changes associated to *ardC*-mediated conjugation. These results show that ArdC is involved in broadening R388 plasmid host range.

## Results

### *ardC* is required for R388 conjugation from *E. coli* to *P. putida*

R388 plasmid is composed by three functional sectors (Fig EV1). One for general maintenance (modules of replication, stable inheritance and establishment) located in the leading region, a sector for Ab^R^ and integration and a third one for conjugation (modules of DNA transfer replication and mating pore formation) [11]. We expected the stable inheritance and establishment region to be required in interspecies conjugation. pSU2007, a Kn^R^ R388 derivative, is transferred with different efficiencies from *E. coli* BW27783-Nx^R^ to other bacteria (Fig EV2). All conjugation experiments were done at the recipient cells optimal growing temperature except for *P. putida* KT2440 to which we observed that the conjugation was more effective at 37 °C. The transfer of pIC10 (R388Δ*kfrA-orf14*), an R388 derivative without the stability and maintenance region, is similar to pSU2007 in all species but in *P. putida* KT2440 where the conjugation frequencies dropped around 1000 times. We have performed the conjugation experiments at both the optimal *E. coli* growing temperature (37 °C) or *P. putida* growing temperature (30 °C). In the stability and maintenance gene region deleted in pIC10 there are 13 genes that code for proteins homologous to some with predicted function of: fertility inhibition in a different system (*kfrA, nuc1, nuc2* and *osa*)[12], hypothetical proteins of unknown function (*orf7, orf8, orf9, orf12* and *orf14*), transcriptional regulators (*ardk* and *klcB*), ssDNA binding protein (*ssB*) and antirestriction (*ardC*). ArdC protein (297 amino acids and 33.2 KDa) has been shown to have an *in vitro* antirestriction function towards Type I and II R-M systems [13]. Thus, we constructed plasmid pLGM25 (R388Δ*ardC*) to check if the effect observed in conjugation with pIC10 could be due to *ardC* gene. This plasmid was introduced into *E. coli* and then conjugated to *E. coli* BW27783-Rf^R^ or *P. putida* KT2440 (Fig 1A). We observed that the absence of *ardC* in the conjugative plasmid pLGM25 reduced conjugation frequency to *P. putida* from 3.8E-02 to 9.0E-05, but not to *E. coli.* Thus, the results observed for pIC10 could be explained to a large extent by *ardC* absence.

**Fig 1.**
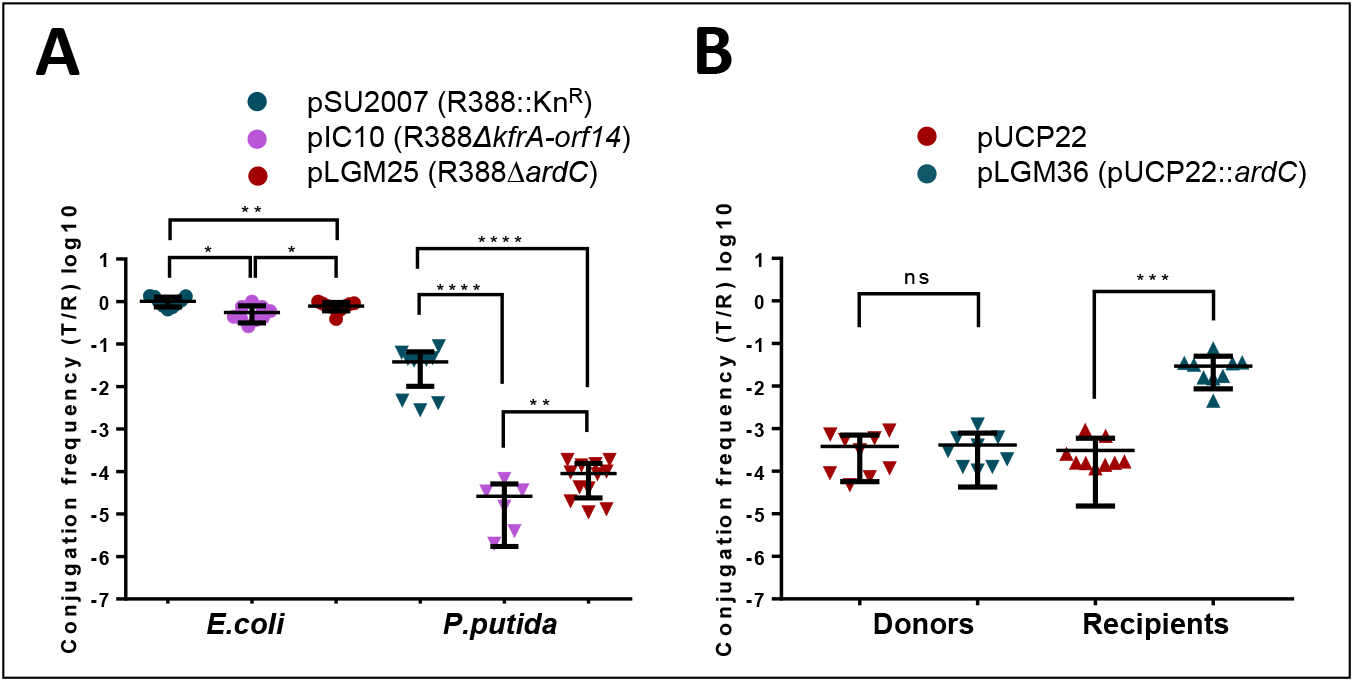
ArdC effect in conjugation. A) Effect of *ardC* and *kfrA-orf14* deletions on plasmid conjugative transfer (1 h at 37 °C) from *E. coli* BW27783-Nx^R^ to *E. Coli* BW27783-Rif^R^ or *P. putida* KT2440. Conjugation frequencies are shown as transconjugants per recipient (T/R). Horizontal bars represent mean ± SD obtained for each dataset of n=6-12 (t-test: * p < 0.1, ** p < 0.01, *** p < 0.001, **** p <0.0001). B) Effect in the conjugation frequency of pLGM25 when expressing *ardC* in donors or recipients. Effect of the presence of plasmid pUCP22 (shown in maroon) or pUCP22∷*ardC* (shown in teal) in donors or in recipients is shown. Conjugation was done for 1 h at 37 °C with 0.1 mM IPTG in the mating mixture. Horizontal bars represent the mean ± SD obtained for each dataset of n=9 (t-test: ** p < 0.01, *** p < 0.001).

### ardC is needed in recipient cells

To check if ArdC is needed in donor or in recipient cells, we have complemented Δ*ardC* pLGM25 plasmid by the overexpression of *ardC* in donor *E. coli (*pLGM25) cells, or in recipient *P. putida* cells. In Fig 1B, we observe how *ardC* does not improve the conjugation frequency when overexpressed in donors. On the other hand, overexpression of *ardC* in recipient cells increased the conjugation frequencies, reaching pSU2007 conjugation levels. Thus, it seems that expression of ArdC is specifically required in recipient cells, and not in donor cells.

### ArdC is a ssDNA-binding protein with a metalloprotease domain

R388 ArdC crystal structure was solved at 2.4 Å resolution using a selenomethionine-derivative protein structure solved by single anomalous dispersion. Using this preliminary structure, the apo ArdC structure was solved at 2.0 Å resolution by molecular replacement (MR). Apo ArdC crystallized in H3 space group containing one molecule per asymmetric unit. Data collection and refinement statistics are given in Table EV1.

ArdC is composed of two structural domains: An N-terminal domain (residues 1-134) and a C-terminal domain (residues 151-297) joined by a long and flexible loop (135-150) (Fig 2A,B), although we could not observe electron density for residues 136-141, in the region connecting both domains. Moreover, electron density was not observed for the N-terminal residues 1-6, C-terminal residues 294-297 or the flexible small loop residues 33-39. The N-terminal domain is composed of three α-helices (α1-α3), a three-stranded β-sheet (β1, β3 and β4) that supports a long and protuberant β-hairpin (β3-β4), a smaller two-stranded antiparallel β-sheet formed by β2 and β5, as well as three 3_10_ helices labelled from η1 to η3 (Fig 2A,B). The C-terminal domain is composed of six α-helices (α4-α9) and three short stranded antiparallel β-sheets (β6-β8) as shown in Fig 2A,B.

**Fig 2.**
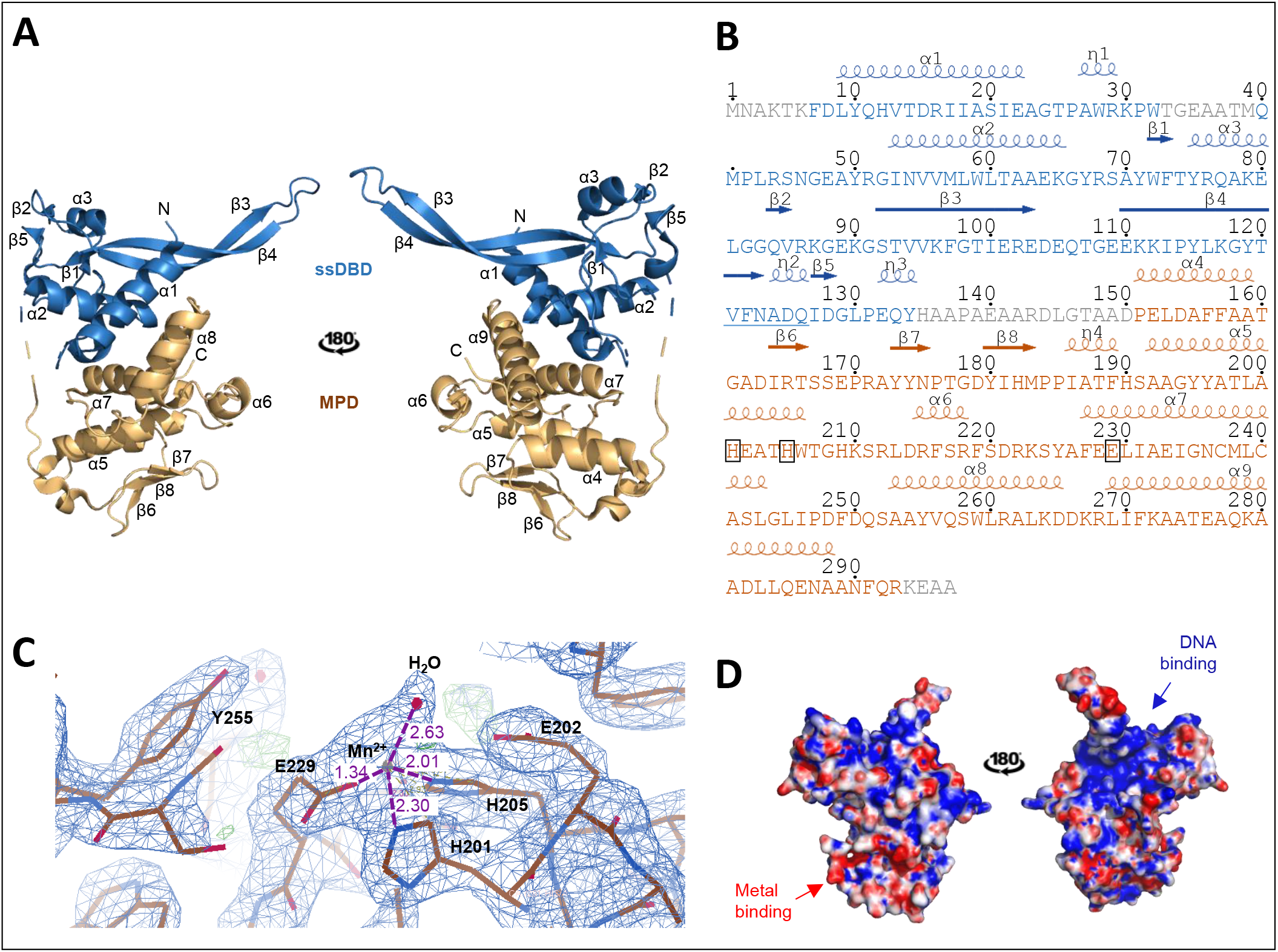
ArdC structure. A) Cartoon representation of two views of ArdC structure. N-terminal ssDNA-binding domain (ssDBD) is shown in blue and C-terminal metalloprotease domain (MPD) in orange. α-helices are labelled from α1 to α9 and β-strands are labelled from β1 to β8. A dashed line schematizes the disordered loop joining both domains. B) ArdC sequence with secondary structure information. ArdC sequence is colored by domains and α-helices and β-strands are labelled as in A). 3_10_ helices are labelled from η1 to η3. The residues involved in metal coordination are framed. The “squiggle” signature proposed by [17] for Rad4 is underlined in blue. C) Electron density of the metal binding site in the ArdC-Mn crystal structure solved at 2.7 Å resolution. Residues and molecules involved in metal coordination (H201, H205, E229 and H_2_O) or activity are labelled. Distance in Å to the metal is shown in purple D) Electrostatic potential surface. Negative surface is colored in red and positive in blue (calculated by APBS tool). The expected binding areas for DNA and metal cofactor are indicated.

ArdC structure was compared to structures deposited in the PDB using the Dali server [14]. The closest structural homologues were the DNA binding metalloproteases Spartan (PDB: 6MDW; Z-score: 6.4) and IrrE (PDB: 3DTE; Z-score: 5.5). Spartan is a protein involved in the cleavage of proteins irreversibly crosslinked to DNA to preserve this way genome stability [15]. IrrE protects *D. radiodurans* from UV radiation DNA damage by proteolysis of a transcriptional regulator, DdrO, involved in SOS response [16].

ArdC N-terminal domain closest structural homolog is in a nucleotide excision repair protein called Rad4 (PDB: 2QSG; Z-score: 4.5), a component of the eukaryotic nucleotide excision repair (NER) pathway. ArdC N-terminal domain possess the V^121^FNADQ^126^ sequence located within a 3_10_ helix (η2) between β4 and β5 (Fig 2B). This region forms a crossover with the β2 to β3 region and creates a sharp twist of the chain known as “squiggle” motif [17]. This squiggle motif in Rad4 is proposed to be responsible of a highly flexible region that could facilitate recognition of DNA sequences. The surface electrostatic map (Fig 2D) reveals a positively charged groove in the region of the N-terminal domain adjacent to the C-terminal domain, suggesting a DNA binding site between both structural domains. By electrophoretic mobility shift assays (EMSA) we have determined that ArdC preferentially binds ssDNA oligonucleotides over dsDNA molecules (Fig EV3A) in accordance to previous results (14). Moreover, binding to partial dsDNA with 5’ or 3’ terminal ssDNA overhangs is preferred over binding to perfectly paired complementary dsDNA duplex (Fig EV3A). We will name ArdC N-terminal domain hereafter ssDNA-binding domain (ssDBD).

Regarding the C-terminal domain, it belongs to the gluzinzin metalloprotease family characterized by the presence of the conserved residues HExxH located on the “active site helix” (α5 in ArdC) and an additional conserved motif (E,H)xx(A,F,T,S,G) located in the contiguous α-helix or “glutamate helix” (α7 in ArdC) [18] (Fig 2A,B). The surface electrostatic map reveals a negatively charged catalytic pocket (Fig 2D). We will name ArdC C-terminal metalloprotease domain MPD.

The two histidines of the active site helix and the glutamic acid of the glutamate helix coordinate a catalytic divalent metal ion, mainly zinc (Fig 2B). However, gluzinzin metalloproteases maintain the catalytic activity with Co^2+^, Mn^2+^ and Ni^2+^ too due to the flexibility of these three metal coordination geometries [18,19]. In order to check the metal used by ArdC we studied the stability of ArdC in the presence of different metal cofactors. ArdC showed an increased thermal stability (assayed by ThermoFluor) in the presence of Ni^2+^, Mn^2+^, and Co^2+^ (ΔT_M_> 5 °C), but not in the presence of Zn^2+^, Ca^2+^, Mg^2+^, Cu^2+^ or Fe^3+^ (Appendix Table S1). Moreover, to know the conformation of the active site when bound to metals, we have crystallized ArdC in the presence of MnCl_2_ (Materials and Methods). ArdC-Mn crystallized in P32 space group containing eight molecules per asymmetric unit and structure was solved at 2.7 Å. We observe how Mn^2+^ is tetrahedrally coordinated by H201, H205, E229 and an H_2_O molecule (Fig 2C). H205 is oriented towards the metal by interaction with the conserved E228 through the non-coordinating nitrogen atom. The E202 of the HE^202^XXH motif orients and acts as a catalytic base for activation of a water molecule that coordinates the metal. The H_2_O molecule could act as a Lewis acid to allow the nucleophilic attack [18]. By analogy with other gluzinzin metalloproteases the conserved ArdC Y255 could stabilize by an hydrogen bond the polypeptide chain to be cleaved [20]. The metalloprotease sub-domain (MPsD) in Spartan shares active center structure with ArdC MPD except that MPsD uses a third histidine instead of a glutamic acid for metal coordination.

It had been proposed that ArdC could avoid ssDNA degradation by HhaI, a type II restriction enzyme able to cleave both ssDNA and dsDNA [13]. According to our structural results, this ArdC DNA protection could be due to ArdC MPD activity targeting the restriction enzyme. To test this hypothesis, we assayed the inhibition of HhaI by ArdC with in the presence of ssDNA M13mp18 (7.2 kb) and Mg^2+^. As observed in Fig EV3BC, ArdC is able to avoid ssDNA cleavage by HhaI but we did not observe HhaI degradation by ArdC.

Since structure defined ArdC as a protease, we tried to find a specific protein target. We purified mutant protein E229A (supposed to be inactive) and used it as prey for co-purification of potential targets in *P. putida* KT2440 cell lysate by pull-down technique. The only protein that eluted at the same time as ArdC was PP_0941, a protein of unknown function similar to the 50S ribosome subunit associated protein YjgA (Appendix Figure S1).

### SOS response is activated in *P. putida* recipient cells by the transfer of an *ardC*-containing plasmid

IrrE, the bacterial closest structural homologue to ArdC, triggers SOS response by cleaving the transcriptional regulator DdrO in an analogous way to the RecA-LexA system [16]. To check if ArdC could have a similar activity on plasmid conjugation, we analyzed by RNA-seq changes in gene expression when an a*rdC*-containing plasmid was transferred from *E. coli* to *P. putida*. As described in materials and methods, we have mixed in a conjugation filter *P. putida* KT2440 with *E. coli* BW27783-Nx^R^ (NP standing for no plasmid), *E. coli* BW27783-Nx^R^ bearing pSU2007 (*ardC+*) or *E. coli* BW27783-Nx^R^ bearing pLGM25 (*ardC* ^−^).

As expected, significant conjugation frequency differences between *ardC* ^+^ and *ardC* ^−^ conditions were observed (Table EV2). RNA-seq results (Datasets EV1, Table EV3-EV4, Appendix Table S2 and Appendix Figure S2) showed: (a) R388 genes involved in conjugation are highly upregulated in the *ardC* ^+^ condition with respect to *ardC* ^−^ conditions (Table EV5, Appendix Table S2 and Appendix Figure S2A). This is consistent with the zygotic induction observed in the recipient cells after conjugation [21] (b) Downregulation of a number of donor *E. coli* genes and pathways involved in flagellar motility, SOS and stress responses and different metabolic pathways in *ardC* ^−^ condition with respect to NP or *ardC* ^+^ conditions (Appendix Table S2, Table EV6 and Appendix Figure S2B). (c) Upregulation of SOS genes in recipient *P. putida* cells when receiving *ardC* ^+^ containing plasmid with respect to NP or *ardC* ^−^ conditions. Differential expression of *P. putida* genes in recipient cells when the ardC-containing plasmid is transferred is shown in Table EV7.

### ArdC metalloprotease activity is not required for conjugation to *P. putida*

Being ArdC a ssDNA binding protein with a metalloprotease domain, we have checked if this proteolytic activity is needed for ArdC activity in conjugation. We mutated *ardC* gene to *ardC_E229A* in pSU2007 to generate plasmid pLGM33. The glutamic acid of the active site E229A, involved in metal coordination, is expected to deactivate the proteolytic center of ArdC, as it occurs in other family members. We conjugated pSU2007 or pLGM33 from *E. coli* to *P. putida*. Surprisingly, pLGM33 conjugation frequency resulted to be like pSU2007, around 0.01 transconjugants per recipient (T/R) (Fig EV4A). In addition, we also tested if ArdC_E229A protein complements pLGM25 (R388 *ΔardC*) plasmid conjugation from *E. coli* to *P. putida* KT2440. The expression of ArdC_E229A in *P. putida* KT2440 recipient cells was able to increase the conjugation frequency of pLGM25 at the same levels than the expression of wt ArdC (Fig EV4B). Thus, ArdC metalloprotease activity is not required for host range broadening activity at least to *P. putida*.

### ArdC counteracts *P. putida* HsdRMS system

To identify the functional target of ArdC, we used a series of *P. putida* mutant strains as recipients. RecA dependent SOS response is activated in *ardC* ^+^ conjugation recipient cells as shown by the RNA-seq experiments (Table EV7). To check the role of this response in conjugation, we first conjugated pSU2007 and pLGM25 to *P. putida* KT2440*ΔrecA*. The frequency of conjugation of pSU2007 to *P. putida* KT2440*ΔrecA* was about 10^−2^ T/R. Thus, RecA dependent SOS response in *P. putida* recipient cells is not essential for conjugation. Moreover, the frequency of conjugation of pLGM25 to *P. putida* KT2440*ΔrecA* was around 10^-4^ T/R, meaning that the absence of *recA* does not enhance conjugation in the absence of *ardC*. In addition, we conjugated to *P. putida* EM42, which carries deletions of several prophages and other accessory genes (*Δ*prophage1 *Δ*prophage4 *Δ*prophage3 *Δ*prophage2 *Δtn7 ΔendA-1 ΔendA-2 ΔhsdRMS Δ*flagellum *Δtn4652*) that could harass the heterologous gene expression (because their association to genetic instability or attributed to the unfruitful usage of metabolic resources). Any of these genes removed from *P. putida* EM42 could avoid the establishment of the plasmids acquired by conjugation. The conjugation frequency towards EM42 strain was not affected by *ardC* deletion (Fig 3) indicating that ArdC could be counteracting the action of the products of one or more of the deleted genes in EM42 strain. To identify the gene or genes responsible for the observed phenotype, we conjugated pSU2007 or pLGM25 from *E. coli* to *P. putida* mutants with deletions in each single gene or group of genes. We could observe that pLGM25 mutant only reached pSU2007 conjugation levels (around 0.1 T/R) in the *P. putida* KT2449*ΔhsdRMS* strain (Fig 3). Similar results were observed at 30 °C (Fig EV5). The *hsdRMS* operon was the main responsible for the effect observed in EM42. Thus, *ardC* is counteracting the effect of the HsdRMS R-M system in the incoming DNA.

**Fig 3.**
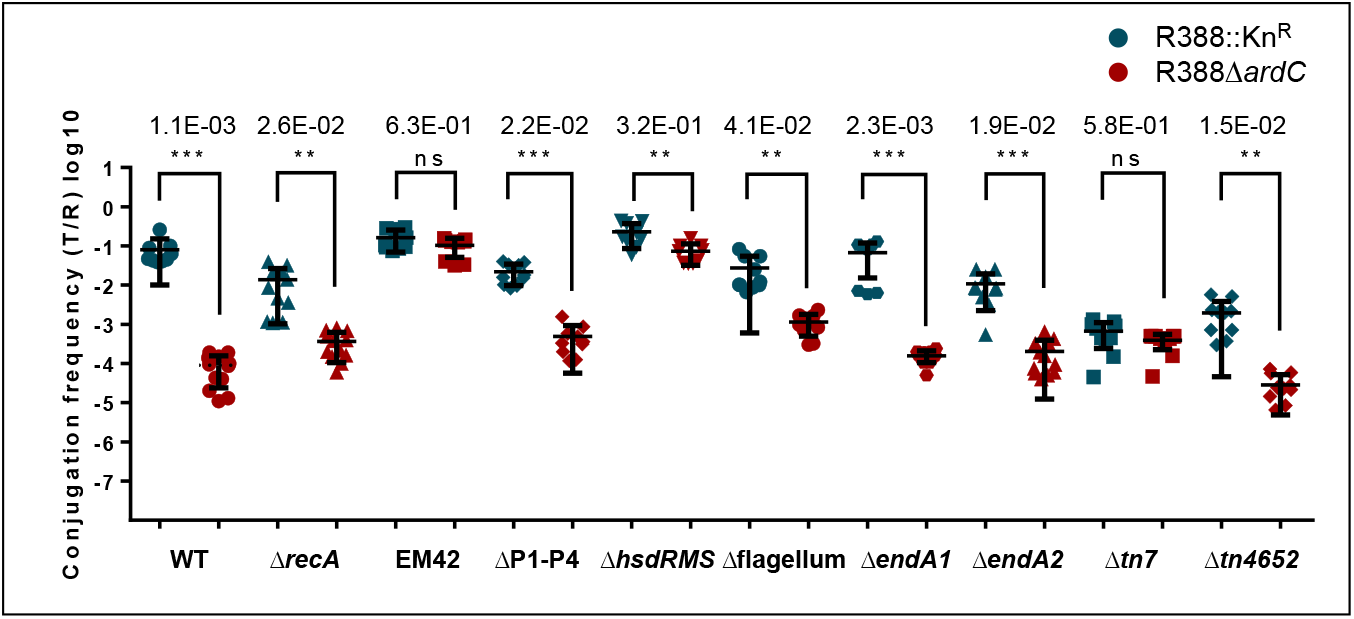
Effect of ArdC on plasmid conjugative transfer from *E. coli* to *P. putida* KT2440 mutants. The conjugation frequencies (T/R) to *P. putida* KT2440 wt strain or to different mutants were obtained after conjugation for 1h at 37 °C. The deleted gene(s) in each strain is shown. EM42 is Δprophage1, Δprophage4, Δprophage3, Δprophage2, Δ*tn7*, Δ*endA-1*, Δ*endA-2*, Δ*hsdRMS*, Δflagellum, and Δ*tn4652*. ΔP1-P4 stands for Δprophage1 Δprophage4 Δprophage3 Δprophage2 strain. Horizontal bars represent the mean ± SD obtained for each dataset of n=8-12 (t-test: ** p < 0.01, *** p < 0.001). Differences in conjugation frequency shown as fold times changes (mean pLGM25/ mean pSU2007) are shown on top for each bacterial strain.

## Discussion

Antibiotic resistance determinants are extensively disseminated by conjugative BHR plasmids. BHR plasmids evolved different strategies to avoid obstacles to their entrance in new recipient cells. In this article, we determined that ArdC protein, produced by the IncW BHR plasmid R388, while dispensable for conjugation from *E. coli* to *E. coli* (Fig EV2), is required for interspecies conjugation from *E. coli* to *P. putida*. ArdC was first studied by [13], who showed an *in vitro* antirestriction function towards Type I and II R-M systems. They observed that ArdC showed 38 % identity with the N-terminal region (about 300 amino acids) of TraC1 primase from RP4 plasmid. Since TraC1 travels to the recipient cell during conjugation bound to the ssDNA that is being transferred (T-strand), they proposed that ArdC could be transferred during conjugation bound to the plasmid T-strand. Besides, they proposed that ArdC protects the incoming DNA from host endonucleases through restriction site occlusion. However, we demonstrate in this work that ArdC role is played directly from its expression in recipient cells, since complementation of pLGM25 (R388Δ*ardC*) with *ardC* in donor cells did not recover pSU2007 conjugation frequencies from *E. coli* to *P. putida*, while the expression of *ardC* in the recipient cells allows high frequency pLGM25 conjugation. Thus, ArdC does not need to travel bound to the ssDNA from donors to recipients, contrary to what was proposed by Belogurov *et al.* [13].

Further information about ArdC activity was obtained by solving the crystal structure of ArdC. Interestingly, ArdC is shown to contain an N-terminal ssDNA binding domain and a C-terminal metalloprotease domain. ArdC is structurally similar to DNA-binding dependent metalloproteases involved in the maintenance of genetic stability such as Spartan or IrrE. Human Spartan protein cleaves DNA-protein crosslinks [15] meanwhile IrrE plays a central regulatory role in DNA protection and repair pathways in response to radiation [16]. ArdC structure differs from other known plasmid-coded antirestriction proteins. ArdA (2W82) is structurally similar to the B-form DNA this way binding type I R-M systems to avoid DNA degradation [22]. KlcA (2KMG) and ArdB (2WJ9) are composed of a single α/β domain inhibiting the endonuclease activity of Type I R-M systems by an indirect mechanism not related to the mimic of DNA structure [23]. According to our solved structure, we expect ArdC to provide a new antirestriction mechanism.

We observed SOS response activation in *P. putida* recipient cells during *ardC* ^+^-mediated conjugation. ArdC could trigger SOS response in recipient cells in a similar way than IrrE triggers SOS response upon radiation damage (by proteolysis of DdrO, a transcriptional regulator involved in SOS response) [24]. However, it has also been described that ssDNA in conjugation activates SOS response [25]. As ArdC_E229A mutant is still able to promote R388 conjugation to *P. putida*, we think that activation of the SOS response in the recipient is the consequence of the ongoing conjugative process and not a direct consequence of the presumed proteolytic activity of ArdC. However, *ardC* ^−^ plasmid pLGM25 should also be entering *P. putida* as ssDNA and thus should activate SOS response, according to results of Baharoglu and Mazel [25]. There is no reason to think that pLGM25 DNA is not being transferred, as it has the complete machinery for channel formation and DNA processing. Furthermore, we observed significant downregulation of genes involved in general bacterial metabolism in donor *E. coli* cells when bearing *ardC* ^−^ plasmid. Donor cells need to make a strong metabolic effort to transfer plasmids [see [21]]. This energy-consuming process should end as soon as the plasmid is established in the recipient cells. However, since pLGM25 cannot be established, we propose that the donor cell keeps pumping ssDNA to the recipient cell. Nevertheless, we did not observe SOS activation in recipient cells as expected for a cell with an abnormal amount of ssDNA [25].

Since ArdC is not required for R388 conjugation to *P. putida* KT2440Δ*hsdRMS*, it is expected to play a role as counteracting Type I R-M system, probably preventing degradation of the transferred DNA. Type I R-M systems attack dsDNA and thus, are not expected to degrade ssDNA during bacterial conjugation. However, it has been reported that the EcoKI R-M system affects the uptake of DNA by conjugation [26] and, ArdC is not expressed until plasmid DNA is in dsDNA shape. Thus, Type I R-M system could be attacking late, once ArdC is generated from the dsDNA plasmid after DNA entrance and establishment in recipient cells. In this respect, the observation that mutation of the ArdC metalloprotease active center does not reduce interspecies conjugation suggests that the metalloprotease activity is not required during conjugative transfer to *P. putida.* The presumed metalloprotease activity is not expected to play a role in HsdRMS activity. Thus, it is tempting to propose that, just by ArdC binding to DNA, the protein interferes with HsdRMS binding and thus interferes with degradation of its target DNA.

In summary, our results indicate a new mechanism of DNA antirestriction played by protein ArdC, by which plasmids increase their conjugation host range. Interfering with ArdC activity could thus provide a new tool to hinder the transmission of antibiotic resistance.

## Materials and Methods

### Culture conditions and antibiotic usage

All bacterial strains (Table EV8) were cultured in Luria-Bertani (LB) medium at 37 °C except *Agrobacterium tumefaciens and Pseudomonas putida* for which the optimal growing temperature was 30 °C. When required, the following antibiotics were used: 100 μg/mL ampicillin, 50 μg/mL kanamycin, 25 μg/mL chloramphenicol, 10 μg/mL gentamycin, 20 μg/mL nalidixic acid, 10 μg/mL tetracycline, 20 μg/mL trimethoprim, 50 μg/mL rifampicin, and 300 μg/mL streptomycin.

### Construction of pET29c derived overexpression vectors

*ardC* coding sequence described by [13] was cloned into pET29c vector (Table EV9) with a C-terminal 6xHis tag by Isothermal assembly [27]. *ardC* was obtained with “ArdC-Nterm” and “ArdC-Cterm” oligonucleotides and vector opened with “pET29CNdeI” and “pET29CXhoI” oligonucleotides (Table EV10) by PCR with Phusion® DNA polymerase (Thermo Scientific). PCR products were digested with DpnI FD (Thermo Scientific) restriction enzyme and incubated in isothermal assembly reaction mixture [27] for 1 h at 50 °C.

This pET29c∷*ardC* construction was mutated by Quick Change site directed mutagenesis method (Adapted from QuickChange II Site-Directed Mutagenesis Kit Protocol) to obtain pET29c∷*ardC_E229A*. We used oligonucleotides *“*ArdC E229A d” and “ArdC E229A r” (Table EV10) and Vent® DNA polymerase. PCR products were digested with DpnI FD (Thermo Scientific) restriction enzyme.

### Construction of pUCP22 derived expression vectors

*ardC* gene (NCBI Reference Sequence: NC_028464.1) was amplified by PCR with oligonucleotides “ardC_rev(Hin)” and “ardC_fwd(Eco)” (Table EV10) and Phusion® DNA polymerase (Thermo Scientific). PCR product and pUCP22 vector (Table EV9) were digested with FastDigest EcoRI and HindIII (Thermo Scientific) restriction enzymes and ligated with T4 DNA ligase (Thermo Scientific).

The resulting pUCP22∷*ardC* plasmid was subjected to Quick Change site-directed mutagenesis done with PfuUltra II Hotstart PCR Master Mix (Agilent Technologies) and oligonucleotides “ArdC E229A d” and “ArdC E229A r” (Table EV10) to generate pUCP22∷*ardC_E229A.*

### Construction of R388Δ*kfrA-orf14* deletion mutant

R388Δ*kfrA-orf14* plasmid (pIC10) was constructed by a modification of the Wanner and Datsenko method [28]. “KfrAKamiK1” and “Orf14KamiK2” oligonucleotides (Table EV10) were used to amplify by PCR with Phusion® DNA polymerase (Thermo Scientific) the Kn^R^ gene (with two added KpnI sites flanking) from pUA66 vector (Table EV9). The 973 bp DNA fragment was extracted from gel, treated with FD DpnI restriction enzyme, and dialyzed against water. The cell strain used for recombination was *E. coli* DY380 (Table EV8). R388 plasmid was introduced by conjugation to this strain and transconjugants (Sm^R^, Tp^R^) were grown at 30 °C o/n. Next day, a 1/50 dilution was done in LB and cells were grown until an OD_600_ of 0.5-0.7. Then, the culture was incubated at 42 °C for 20 min with shaking to induce the recombineering system. After this time, cells were place on ice for 20 min and made electrocompetent according to standard molecular biology protocols [29]. For transformation, 100 ng of the PCR product were used. Electroporation was performed at 2.5 keV and time constant (3-5 ms) in a Micropulser™ electroporator (Bio-Rad), cells were immediately recovered in 1 mL of sterile LB prewarmed at 30 °C. Then, cells were let at 30 °C for 2 h. Then, cells were plated in Kn and incubated at 30 °C o/n. Colony checking PCRs were done with Taq polymerase to verify the substitution. Plasmid from some positive colonies were conjugated to DH5α cells by conjugation for 1 h at 30°C and plated on Kn Nx LB agar plates. Another checking colony PCR was done to confirm that we had the mutant plasmid isolated.

### Construction of R388Δ*ardC* deletion mutant

R388Δ*ardC* plasmid (pLGM25) was constructed by a modification of the Wanner and Datsenko method [28]. “N_Kn_promoter_Wanner” and “C_Kn_Wanner” oligonucleotides (Table EV10) were used to amplify by PCR with Vent® polymerase the Kn^R^ cassette with its promoter from pET29c vector (Table EV9). The 1227 bp DNA fragment was extracted from gel, treated with FD DpnI restriction enzyme, and dialyzed against water. The cell strain used for recombination was *E. coli* TB10 (Table EV8). R388 plasmid was introduced by conjugation to this strain and transconjugants (Tc^R^ and Tp^R^) were grown at 30 °C o/n. Next day, a 1/70 dilution was done in LB and cells were grown until an OD_600_ of 0.5. Then, the culture was incubated at 42 °C for 15 min with shaking to induce the recombineering system. After this time, cells were made electrocompetent according to standard molecular biology protocols [29]. For transformation, 100 ng of the PCR product were used. Electroporation was performed at 2.5 keV and time constant (3-5 ms) in a Micropulser™ electroporator (Bio-Rad), cells were immediately recovered in 1 mL of sterile LB prewarmed at 30 °C. Then, cells were let at 30 °C for 3 h. Then, cells were plated in Tc and Kn at half the normal antibiotic concentration. After one day, colonies were restricken in a plate with the normal antibiotic concentration. Colony checking PCRs were done with Taq polymerase with oligonucleotides “Up”, “Down”, “Middle_Up” and “Middle_Down” (Table EV10) to verify the substitution. Plasmid DNA from some of the positive colonies was extracted and introduced in DH5α cells by electroporation to make sure that only one type of plasmid (mutated or WT) entered each cell. Another checking colony PCR was done for the selected mutated colonies with oligonucleotides “ArdC-Cterm” and “ArdC-Nterm” (Table EV10) to make sure that they do not amplify any fragment and thus, confirm that we had the mutant plasmid isolated.

### Conjugation assays

Donor and recipients cell cultures were grown o/n at their optimal growing temperature with shaking (120 rpm) in the presence of the selecting antibiotics. After OD_600_ measurement, the needed volumes to have an OD_600_ (D: R) = 0.6:0.6 in 1 mL were mixed and washed twice by resuspension in 1 mL LB and centrifugation to remove the antibiotics. After the last centrifugation, the cell mixture was resuspended in 30 μL of LB. The conjugations were done in solid LB agar plates previously incubated at the mating temperature with a 0.22 μm pore size cellulose acetate filter of 25 mm of diameter (Sartorius Stedim). The 30 μL of conjugation mixture were placed over the filter and kept at the conjugating temperature for the desired time. If not specified, the standard conditions were 1 h at 37 °C. After this time, the filters were removed with sterile tweezers and introduced in 1 mL LB, where the cells were resuspended by vortexing for a few seconds to stop the conjugation. This tube was considered dilution zero. 1/10 serial dilutions were done and 10 μL drops were plated in LB agar plates with the appropriate selecting antibiotics for donors, recipients and transconjugants. Conjugation frequencies were obtained by dividing transconjugants per recipients (T/R). For conjugations in the presence of pUCP22-derived plasmids, IPTG was added to the conjugation mixture to a 0.1 mM IPTG final concentration.

### Statistical analysis

Means and standard deviations, as well as statistical tests were calculated with GraphPad Prism^®^ (v 7.04) biostatistics software.

### Electrophoretic mobility shift assay (EMSA)

The binding ability of ArdC protein with ssDNA, dsDNA and dsDNA with ssDNA overhangs were tested by electrophoresis mobility shift assay (EMSA). 6FAM-labelled oligonucleotide Fluor-T87I2 (45b) was incubated alone or with T87I1 (complementing 45b), Mid1 (complementing 5’ terminal 18b) or Mid2 (complementing 3’ terminal 27b) in buffer containing 50mM TrisHCl (pH 7.5) and 1mM EDTA for 5 min at 95 °C and let them slowly cold down o/n to room temperature. Ten microliters reaction mixture containing 50 nM DNA was incubated with various concentrations of ArdC (0, 125 nM, 250 nM, 500 nM and 1 μM) in a reaction buffer [50 mM NaCl, 25 mM Tris-HCl (pH 7.5), 0.5 mM EDTA] at room temperature for 30 min. DNA-protein complexes were analyzed using non-denaturing polyacrylamide gel electrophoresis 10 % (29:1) in cold Tris Borate EDTA (TBE) buffer 1x. Gels were run at 100 V, during 75 min and analyzed using a Fujifilm fluorescent image analyzer Fla-5100. Experiments were repeated three times.

### Transcriptomic analysis

Mixtures of *E. coli* BW27783-Nx^R^ (bearing pSU2007, pLGM25 or no plasmid) with *P. putida* KT2440 were carried out by the already described conjugation assay with the following modifications: A ratio of five donor cells per recipient was chosen to make sure that all the recipient cells could be in contact with a donor to start the conjugation process.

The traditional protocol of conjugation was shortened to 30 min in an attempt to obtain the RNA synthesized in the recipient cell during the conjugation process.

Harvested cells obtained from the conjugation filter were treated with two volumes of RNAprotect® Bacteria Reagent (Qiagen). Cells were centrifuged, snap-frozen, and stored at -80 °C. Cells were lysed with 5 μg lysozyme (Sigma) and 50 ng proteinase K (Roche). After cell lysis, total RNA was extracted with RNeasy® Mini Kit (Qiagen) and treated with RNase-free DNase (Qiagen) in column for DNA removal. Ambion® TURBO DNA-*free*™ DNase Treatment was also applied for a better DNA removal. RNA integrity and quality were validated by the Agilent RNA ScreenTape assay. The RNA integrity number equivalent (RIN^e^) was assured to be above 8 to use the isolated RNA in the RNA-seq experiment.

Transcriptome libraries were prepared by Macrogen (Seoul, Korea) with Ribo-Zero rRNA Removal Kit and TruSeq® Stranded mRNA sample preparation kit (Illumina) by following the Low Sample LS protocol. Libraries were sequenced by Macrogen on the Illumina HiSeq 4000 platform. The transcriptome libraries were paired-end sequenced with 100-bp reads.

Raw reads in FASTQ format were quality analyzed with FastQC [30]. For mapping the reads, sequences of R388 (NCBI Accession number NC_028464.1), *E. coli* str. K-12 substr. MG1655 (U00096.3) and *P. putida* KT2440 (AE015451.2) were used as genome template. The alignment of reads was done by each side independently with Bowtie2 software [31]. Artemis program [32] was used to visualize the alignment and do the RPKM (reads per kilobase and million mapped reads) calculations. Genes with less than 10 RPKMs in all experimental conditions were removed from the analysis. DAVID online tool v6.8 [33] was used to test for gene ontology enrichment among the list of differentially expressed genes to do a functional classification.

### Protein expression and purification

Cloned constructs pLGM21 (pET29c∷*ardC*) and pLGM28 (pET29c∷*ardC_E229A*) were transformed into electrocompetent *Escherichia coli* BL21 (DE3) cells (Table EV8). Transformed cells were grown in 1L LB medium, in the presence of kanamycin, at 37 °C, with shaking, to an optical density of 0.5–0.6. The temperature was reduced to 18 °C and protein expression was induced with Isopropyl β-D-thiogalactoside (IPTG) to a final concentration of 0.5 mM. Cells were allowed to grow for 16 h. The resulting growths were spun at 5,000 rpm at 4 °C for 15 min and stored at −20 °C. Pellet was resuspended in buffer A (500 mM NaCl, 20 mM imidazole, 100 mM Tris-HCl pH 7.5) supplemented with protease inhibitor phenylmethylsulfonyl fluoride (PMSF) 1 % (v/v). The slurry was sonicated in a *Labsonic 2000 (B. Braun)* equipment at 50 % of potency for 3 cycles of 1.5 min at intervals of 1 min on ice. The lysed cells were then ultra-centrifuged at 40,000 rpm for 15 min at 4 °C. The supernatant was filtered and purified over a 5 mL nickel HisTrap™ HP (GE Healthcare) column eluting by a lineal gradient between buffer A and buffer B (300 mM NaCl, 500 mM imidazole, 100 mM Tris-HCl pH 7.5). ArdC containing fractions were pooled and diluted to a final NaCl concentration of 200 mM. The resulting protein was then further purified over an affinity chromatographic HiTrap® Heparin HP (GE Healthcare) equilibrated with buffer C (100 mM Tris-HCl pH 7.5, 200 mM NaCl). Elution of bound proteins was done by a lineal gradient between buffer C and D (100 mM Tris-HCl pH 7.5, 1 M NaCl). An additional step of size exclusion chromatography was done through a gel filtration Superdex™ S75 column 10/300 GL (GE Healthcare) with buffer E (100mM Tris-HCl pH 7.5, 1 mM EDTA, 300 mM NaCl).

In order to crystallize ArdC with a metal cofactor, all the lysis and the two first purification steps were done as described but with an additional 1 mM MnCl_2_ final concentration in all buffers. Preparation of selenomethionine (SeMet)-labelled ArdC was also carried out as described above but using strain *E. coli* β834 (DE3) and minimal medium (SelenoMet™ Medium Base + SelenoMet™ Nutrient Mix) supplemented with SelenoMethionine Solution (Molecular Dimensions) as indicated by the manufacturer.

### Protein crystallization and structure determination

Crystals of ArdC and ArdC-SeMet were obtained using the sitting-drop vapor-diffusion method at 22 °C by mixing 1.5 μL protein at 20 mg/mL concentration in 20 mM Tris-HCl, 50 mM NaCl, 1 mM EDTA buffer with an equal volume of the reservoir solution containing 0.1 M HEPES pH 7.5; 10 % w/v polyethylene glycol 6,000 and 5 % v/v (+/−)-2-methyl-2,4-pentanediol. 2-methyl-2 4-pentanediol (10-20 % v/v) was added as the cryoprotectant before diffraction experiments. ArdC-Mn crystallized at 12 mg/mL in 25 % v/v ethylene glycol and crystals were cryoprotected with an additional 15 % glycerol.

The crystals were flash frozen in liquid nitrogen for data collection at 105 K and 12.66 KeV by rotating the single frozen crystals in Δφ= 0.25° steps through 180°-360°. For the single SeMet crystals, a fluorescence scan was performed at the selenium absorption edge to confirm that the selenomethionine substitution was successful, and data was collected at 0.9793Å, the wavelength corresponding to the heavy atom absorption maximum. Datasets were obtained at beamline XALOC at the ALBA Synchrotron Radiation Facility (Barcelona, Spain) with a Dectris PILATUS3 6M Pixel detector. Diffraction images were processed using iMosflm [34] and Scala [35] as part of the CCP4 package [36]. For solving the phase problem, single anomalous dispersion (SAD) method was used thanks to the selenium introduced in the protein. For improving the resolution, data took from native protein crystals was used, solving the phase problem by molecular replacement (MR) with the structure obtained by SAD and using the program MolRep (CCP4). Refinement of the initial model was performed through several cycles by Phenix refine [37] until appropriate R factors are reached. Final manual modelling was done in COOT [38]. For ArdC-Mn structure, MR was also used.

### Mass spectrometry analysis for protein identification

To identify the putative ArdC protease target obtained by the pull-down assay, a protein identification assay was done by Liquid chromatography-tandem mass spectrometry (LC-MS/MS). Selected protein band was subjected to in-gel tryptic digestion according to [39], with minor modifications. Gel pieces were swollen in digestion buffer containing 50 mM NH_4_HCO_3_ and 12.5 ng/μL proteomics grade trypsin (Roche, Basel, Switzerland), and the digestion processed at 37 °C o/n. The supernatant was recovered and peptides were extracted twice: first, with 25 mM NH_4_HCO_3_ and acetonitrile (ACN), and then with 0.1% (v/v) trifluoroacetic acid and ACN. The recovered supernatants and extracted peptides were pooled, dried in a SpeedVac (ThermoElectron, Waltham, MA) dissolved in 10 μL of 0.1 % (v/v) formic acid (FA) and sonicated for 5 min. LC-MS/MS spectra were obtained using a SYNAPT HDMS mass spectrometer (Waters, Milford, MA) interfaced with a nanoAcquity UPLC System (Waters). An aliquot (8 μL) of each sample was loaded onto a Symmetry 300 C18, 180 μm × 20 mm precolumn (Waters) and washed with 0.1 % (v/v) FA for 3 min at a flow rate of 5 μL/min. The precolumn was connected to a BEH130 C18, 75 μm × 200 mm, 1.7 μm (Waters), equilibrated in 3 % (v/v) ACN and 0.1 % (v/v) FA. Peptides were eluted with a 30 min linear gradient of 3−60 % (v/v) ACN directly onto a homemade nano-electrospray capillary tip. Capillary voltage was set to 3,500 V and data-dependent MS/MS acquisitions performed on precursors with charge states of 2, 3, or 4 over a survey m/z range of 350−1990. Raw files were processed with VEMS [40] and searched against the NCBI non-redundant (nr) database restricted to Proteobacteria (version 20171205, 49911253 sequences) using the online MASCOT server (Matrix Science Ltd., London; http://www.matrixscience.com). Protein identification was carried out by adopting the carbamidomethylation of Cys as fixed modification and the oxidation of Met as variable modification. Up to one missed cleavage site was allowed, and values of 50 ppm and 0.1 Da were set for peptide and fragment mass tolerances, respectively. Mass spectrometry analysis were performed in the Proteomics Core Facility-SGIKER (member of ProteoRed-ISCIII) at the University of the Basque Country, UPV/EHU.

### DNA-binding and protection assays

The assay was done under non-denaturing conditions to see DNA binding and retardation in parallel to under denaturing and proteolytic conditions: M13mp18 ssDNA (7.2 Kb, 5.5 nM final concentration) was incubated with increasing concentrations of ArdC for 10 min at RT in a total volume of 20 μL binding buffer: 10 mM Tris-HCl, 10 mM NaCl and 10 mM MgCl_2_. Then, 7 U of HhaI were added and incubated for 20 min at 37 °C. Afterwards, 2.5 μL of DNA loading buffer were added to 10 μL of the sample. Samples were subjected to electrophoresis on a 1 % agarose gel with SYBER safe for 30 min at 120 V. 1.5 μL of proteinase K at 20 mg/mL and 1 μL SDS 10 % were added to the remaining 10 μL of the sample and the mixture was incubated for another 20 min at 37 °C. Reactions were mixed with 2.5 μL DNA loading buffer and electrophoresed in a 1 % agarose gel with SYBER safe for 30 min at 120V.

### Proteolytic activity assay

ArdC proteolytic activity was analyzed using a modification of the method described by [41] for the study of IrrE metalloprotease. ArdC at a final concentration of 8 μM in 20 μL was incubated in buffer P (15 mM Tris-HCl pH 7.5, 15 mM NaCl and 15 mM of MgCl_2_ or 15 mM EDTA) and 27.5 nM M13 ssDNA for 10 min at RT. Then 70 U of HhaI were added and incubated for 20 min at 37 °C. Reactions were stopped by the addition of 20 μL of protein loading buffer 2x (400 mM Tris-HCl pH 6.8, 4 % SDS, 30 % glycerol and 0.04 % bromophenol blue) and boiled for 5 min. A 12 % SDS-PAGE was performed for 60 min at 180 V.

### Site-directed MAGE *in vivo* mutagenesis method

To perform point mutation in pSU2007 and construct pLGM33, we used the non-automated version of the MAGE (Multiplex Automated Genome Engineering) method described by [42]. ArdC_E229A_MAGE” (Table EV10), a 90 base oligonucleotide containing the mutation in the middle and two phosphorothioate (PS) bonds in the 5’ end was designed. The phosphorothioate bond substitutes a sulphur atom for a non-bridging oxygen in the phosphate backbone of the primer. This modification in the internucleotide bond makes the primer resistant to exonuclease degradation. EcMR2Δ*mutS E. coli* strain (Table EV8) was used. These cells were cultured at 30 °C when recombination was not needed. pSU2007 plasmid was introduced by conjugation into this strain and transconjugants (Kn^R^ and Rif^R^) were grown at 30 °C o/n. Next day, a 1/40 dilution was done in LB and cells were grown until an OD_600_ of 0.5. Then culture was incubated at 42 °C for 15 min with shaking to induce the recombineering system and after this time cells were made electrocompetent according to standard molecular biology protocols [29] except for the last wash, when cells were resuspended in 50 μL of a 1 μM oligonucleotide suspension so the mixture is ready for electroporation. Once recovered in 1 mL LB, cultures were grown at 30 °C. After 2 h, 50 μL were plated in Kn Rif plates and the rest of the volume was grown o/n labelled as cycle #1 until next day when the protocol was repeated. When needed, stocks were saved at −80 °C in 25 % glycerol for further analysis. After 10 cycles, plasmid extraction from some colonies was done and the PCR product obtained with Phusion® polymerase and oligonucleotides “Up” and “Down”, was sent to sequence with “Down” oligonucleotide. The colony that gave an overlapping pick at the sequencing pane in the mutagenic position was further analyzed by reelectroporating in DH5α and sequencing the PCR fragments from some of the colonies until a clean mutant pick was obtained for the desired position.

### Thermal stability assay based on fluorescence

20 μL samples containing ArdC at 2.5 μM in 100 mM Tris-HCl pH 7.5 and 500 mM NaCl buffer containing 1mM of EDTA or 1mM of the metal to be analyzed and the SYPRO® Orange (Invitrogen) non-polar dye at a 2x final concentration were evaluated in a StepOnePlus™ Real-Time PCR System (Thermo Fisher). For measuring SYPRO® Orange (excitation: 470 nm/ Emission: 570 nm), filter for NED™ dye was used, with excitation at 546 nm and emission of 575 nm. Temperature was raised from 25 °C to 85 °C at 0.5 °C per minute, measuring the fluorescence every 0.5 °C. T_M_ was determined as the maximum of the fluorescence versus temperature variation (dF/dT). The experiments were done by duplicate.

## Data availability

The datasets and computer code produced in this study are available in the following databases:

Coordinates and structure factors of ArdC X-ray crystal structure: PDB 6I89
Coordinates and structure factors of ArdC-Mn X-ray crystal structure: PDB 6SNA

## Acknowledgments

We thank Victor de Lorenzo’s group at the Centro Nacional de Biotecnología (CNB) for sharing *P. Putida* strains. We also thank Robert E.W. Hancock’s group at the University of British Columbia (UBC) for pUCP22 vector and financial and technical support to construct pUCP22∷*ardC* plasmid. Structural experiments were performed at XALOC beamline at ALBA Synchrotron with the collaboration of ALBA staff. This research was supported by grant BFU2014-55534-C2 of the Spanish Ministry of Economy, Industry and Competitiveness to F.dlC. and G.M. L.G-M. was recipient of FPU014/06013 fellowship by the Spanish Ministry of Education, Culture and Sports.

## Author contributions

F.dlC. and G.M. designed research; L.G-M. and I.dC. performed research; L.G-M., F.dlC and G.M analysed data; and L.G-M., F.dlC. and G.M. wrote the paper.

## Expanded View Figure Legends

**Figure EV1. Genetic map of R388 plasmid**. The figure shows the genetic organization of the plasmid divided in functional modules and three different sectors: conjugation shadowed in orange, general maintenance in blue and Ab^R^ and integration in grey. Region deleted in pLGM25 (shown in maroon) and pIC10 (shown in purple) are also shown. Adapted from [11].

**Figure EV2. Effect of** *kfrA-orf14* **region on R388 plasmid conjugative transfer from** *E. coli* **to different bacteria**. Conjugation frequencies per recipient (T/R) for R388 or pSU2007 are shown in teal and for pIC10 in purple. Conjugations were performed as described in Materials and Methods at 37 °C except for *P. putida* and *A. tumefaciens* (done at 30 °C) for 1 h except for *A. baumanii* and *V. cholerae* (done for 4 h). R388 was used in conjugations towards *E. coli, S. typhimurium, K. pneumoniae*, meanwhile pSU2007 was used in conjugations towards the rest of the strains. Donor *E. coli* BW27783-Rif^R^ cells were used towards *E. coli*, *S. typhimurium, and K. pneumoniae* meanwhile *E. coli* BW27783-Nx^R^ cells were employed in conjugations towards *the rest of the strains.* Horizontal bars represent the mean ± SEM of n=9-20 (t-test: * p < 0.1, ** p < 0.01, *** p < 0.001). Differences in conjugation frequency shown as fold times changes (mean pIC10/ mean R388 or pSU2007) are shown on top for each bacterial strain.

**Figure EV3. Retardation and protection of DNA by ArdC from degradation by HhaI**. A) ArdC DNA-binding preferences as assessed by EMSA. ArdC binding of a 6FAM-labelled 45 bases ssDNA oligonucleotide (Fluor-T87I2), a perfectly paired complementary 45bp dsDNA duplex (Fluor-T87I2 + T87I1) and two partial dsDNA with 5’ or 3’ terminal ssDNA overhangs (Fluor-T87I2 + Mid1) and (Fluor-T87I2 + Mid2) respectively was performed at increasing concentrations of ArdC (0, 125 nM, 250 nM, 500 nM and 1 μM) as described in Materials and Methods. Protein-DNA complexes were resolved by native 10 % polyacrylamide gels and visualized via fluorescent image analyzer. B) Left: Agarose gel showing ssDNA retardation under non-denaturing conditions. Lane 1: ssDNA (5.5 nM) in the presence of MgCl_2_. The vast majority of the molecules of M13mp18 ssDNA are circular (upper band), although some of them are present in the linear form (lower band). Lane 2: ssDNA and HhaI (7 U) in the presence of MgCl_2_. Lane 3-7: ssDNA and HhaI at increasing concentrations of ArdC (0.95 μM (3), 1.9 μM (4), 3.8 μM (5), 5.7 μM (6) and 7.8 μM (7)) in the presence of MgCl_2_. Right: Agarose gel showing ssDNA protection from HhaI proteolysis under denaturing conditions (proteinase K and SDS added). Lane content as in A). C) SDS-PAGE gel showing proteolytic activity of ArdC preincubated with ssDNA (M13mp18) to HhaI. Lane 1: ArdC (8 μM) in the presence of MgCl_2._ Lane 2: HhaI (70 U) in the presence of MgCl_2_. Lane 3: ArdC and HhaI in the presence of MgCl_2_. Lane 4-5: ArdC and HhaI in the presence of ssDNA (M13, 27.5 nM) with MgCl_2_ (4) or EDTA (5).

**Figure EV4. Effect in the conjugation frequency when ArdC E229 residue is mutated to A. A)** Conjugation was done for 1 h at 37 °C from *E. coli* BW27783-Nx^R^ bearing pSU2007 (shown in teal) or pLGM33 (pSU2007_*ardC_E229A*, shown in orange) to *P. putida* KT2440. Conjugation was done for 1 h at 37 °C. Horizontal bars represent the mean ± SD of n=3 observations. **B)** Complementation of pLGM25 in *E. coli* BW27783 donor cells was done in *P. putida* KT2440 recipient cells bearing pUCP22∷*ardC* (shown in teal) or pUCP22∷ *ardC_E229A* (shown in orange). Conjugation was done for 1 h at 37 °C with 0.1 mM IPTG added to the mating mixture. Horizontal bars represent the mean ± SD obtained for each dataset of n=9.

**Figure EV5. Effect of ArdC on plasmid conjugative transfer from** *E. coli* **to** *P. putida* **wt and mutants at 30 °C**. The conjugation frequencies per recipient (T/R) into *P. putida* KT2440 WT strain or into different mutants of *P. putida* KT2440 are shown. Conjugation was done for 1 h at 30 °C. Horizontal bars represent the mean ± SD obtained for each dataset of n=9 (t-test: ** p < 0.01).

